# Exploring Approximate Bayesian Computation for inferring recent demographic history with genomic markers in non-model species

**DOI:** 10.1101/252650

**Authors:** Joane S. Elleouet, Sally N. Aitken

## Abstract

Approximate Bayesian computation (ABC) is widely used to infer demographic history of populations and species using DNA markers. Genomic markers can now be developed for non-model species using reduced representation library (RRL) sequencing methods that select a fraction of the genome using targeted sequence capture or restriction enzymes (genotyping-by-sequencing, GBS). We explored the influence of marker number and length, knowledge of gametic phase, and tradeoffs between sample size and sequencing depth on the quality of demographic inferences performed with ABC. We focused on 2-population models of recent spatial expansion with varying numbers of unknown parameters. Performing ABC on simulated datasets with known parameter values, we found that the timing of a recent spatial expansion event could be precisely estimated in a 3-parameter model. Taking into account uncertainty in parameters such as initial population size and migration rate collectively decreased the precision of inferences dramatically. Phasing haplotypes did not improve results, regardless of sequence length. Numerous short sequences were as valuable as fewer, longer sequences, and performed best when a large sample size was sequenced at low individual depth, even when sequencing errors were added. ABC results were similar to results obtained with an alternative method based on the site frequency spectrum (SFS) when performed with unphased GBS-type markers. We conclude that unphased GBS-type datasets can be sufficient to precisely infer simple demographic models, and discuss possible improvements for the use of ABC with genomic data.

## Introduction

Patterns of DNA variation among individuals are commonly used to unravel events in the history of populations, such as demographic expansion, population splits, and admixture. Rapid progress in sequencing technologies at the start of the 21^st^ century has allowed the inference of increasingly complex demographic models, by using increasingly complete genomic datasets. However, this increase in amount of data and complexity of demographic scenarios necessitates new statistical methods for analysis and inference. Tackling large genetic datasets with inherent errors and uncertainties requires sophisticated techniques for marker development. In parallel, inferring complex historic demographic scenarios with several populations and numerous demographic parameters necessitates efficient algorithms to provide accurate parameter estimates and model validation measures. Reviews and improvements of methods have recently emerged (Schraiber & Akey, 2015), illustrating the fast pace of change in the field of statistical genetics. However, the efficiency of inference methods for different types of demographic models as well as effects of completeness of genomic datasets need to be understood to ensure quality and accuracy of inferences.

### Demographic inference in natural populations of non-model organisms

In less than 30 years, human demographic inference has taken a leap, evolving from the evidence for a single African origin of all humans using a few non-recombining mitochondrial markers (Cann et al., 1987), to the inference of highly complex demographic scenarios using whole genomes (Harris & Nielsen, 2013). Although there is still room for improvement in demographic inference of human populations (Schraiber & Akey, 2015), human genomics is at the leading edge of inference from DNA data. Unfortunately, the state-of-the-art statistical inference techniques applied to human data are currently out of reach for studies of natural populations of non-model organisms. Knowledge from demographic inference of these species is, however, crucial: it is often the most efficient way to determine how to manage invasive species (Benazzo et al., 2015; Guillemaud et al., 2010), to conserve endangered species or ecosystems (Chan et al., 2014; Dussex et al., 2014; Lopez et al., 2006; Quéméré et al., 2012), and to predict the future distribution and abundance of widespread species that are of economical or ecological importance (Holliday et al., 2010; Zinck & Rajora, 2016). The good news is the genomic revolution has reached non-model organisms, creating a spectrum of levels of genetic knowledge across a broad range of taxa. Using a few microsatellites or moderate-sized panels of resequenced SNPs is still common practice (Y. Li et al., 2010; Zinck & Rajora, 2016), but most current studies of non-model species now use genomic methods to extract markers for inference. In recent years, sequencing whole genomes of non-model species has become feasible in some organisms with small genomes (Boitard et al., 2016; Liu et al., 2014) and has allowed the inference of detailed demographic models using Approximate Bayesian Computation (ABC) or Pairwise Sequential Markovian Coalescent (PSMC) (Nadachowska-Brzyska et al., 2013). For organisms with larger genomes or for studies with lower data requirements, reduced-representation library (RRL) sequencing, through either targeted capture or restriction enzymes, is widely applied (Davey et al., 2011). RRL techniques involving restriction enzymes (commonly referred to as RADseq or genotyping-by-sequencing, GBS) output a large number of short sequences (100bp, or longer with paired-end sequencing) from across the genome and have proven useful in population genetics studies and inference involving maximum likelihood methods based on the site frequency spectrum (SFS) or ABC methods (Narum et al., 2013). Most recently, the number of published drafts of whole genomes for non-model species has increased dramatically, granting access to longer sequences through the second category of genomic markers: targeted enrichment. This approach allows the use of linkage information for population genetics inference (Li & Jakobsson, 2012).

### Approximate Bayesian Computation and other approaches

In this paper, our aim is to explore ABC for datasets obtained from reduced-representation library sequencing in non-model organisms. We also compare the results obtained with those from a SFS approach based on approximation of the composite likelihood (Excoffier & Foll, 2011). We chose to explore ABC because of its versatility: It accommodates a wide spectrum of demographic models and dataset types. Although it was originally developed for inferences in evolutionary biology, the statistical framework of ABC has been extended to a variety of disciplines, from cell biochemistry and epidemiology to neural networks, extending beyond the realm of biology into meteorology, astrophysics (Weyant et al., 2013) and computer sciences (Condon & Cukier, 2016). ABC has been reviewed in a number of publications and its algorithms and techniques are being refined constantly (Bertorelle et al., 2010; Csilléry et al., 2010; Lintusaari et al., 2016; Marin et al., 2012; Sunnaker et al., 2013). For applications in demographic inference using genetic data, the general ABC method involves the following steps. First, a large number of datasets are simulated under a specific demographic model using the coalescent (Kingman, 1982). Parameters used for simulations are drawn from prior distributions that are pre-defined by the user. The simulated datasets are then compared to the observed dataset through calculation of summary statistics. Finally, simulated datasets with the closest vector of statistics to the vector of observed summary statistics are selected. A regression adjustment based on the local relationship between statistics and parameters is then usually performed to approximate the posterior distribution of each model parameter from the parameter values of selected simulations. ABC is suitable when inferring models for which the likelihood function is intractable, as it relies on approximating the likelihood function using a large number of simulations. However, each one of the numerous steps in the implementation of ABC requires users to make empirical decisions. There is particularly a need to improve our understanding of the relationship between the type of markers obtained to build genetic datasets and the way genetic data is subsequently summarized on its power to tease apart demographic models and produce accurate parameter estimates.

### Previous work exploring ABC

The need to test the inference power of datasets for demographic models of interest has been recognized in recent years, both in terms of model selection and parameter estimation. Robert et al. (2011) warned against the use of insufficient summary statistics in ABC model choice, opening the door to improved methods for model testing and the associated choice of summary statistics (Marin et al., 2014; Prangle et al., 2013). Among theoretical results and general guidelines, Marin et al. (2014) suggested the use of different sets of summary statistics for estimation and model selection. Several studies show the use of preliminary simulations testing parameter estimation and model choice with different number and length of markers and number of individuals (Sousa et al., 2012; Stocks et al., 2014), type of molecular markers (Cabrera & Palsbøll, 2017) and choice of summary statistics and models considered (Benazzo et al., 2015; Guillemaud et al., 2010; Li & Jakobsson, 2012; Sousa et al., 2012; Stocks et al., 2014). As most scientists have switched to using genome-wide data, there is a need to expand this set of simulation studies to test and understand the power of different types of genomic data. As part of such an effort, Li & Jakobsson (2012) simulated large, phased genomic datasets comparable to human genomic datasets at the time. Under 2-population split models, they found that ABC produces accurate estimates for most but not all parameters and concluded ABC is well suited to large genomic datasets summarized with LD-based statistics. Robinson et al. (2014) tested the effects of the number and length of unphased genomic sequences and compared them to the effect of the number of individuals sequenced for the inference of three-population admixture models. They found that increasing the number and length of sequences was more beneficial than increasing sample size. Shafer et al. (2015) investigated the power of ABC on short diploid sequences obtained by GBS. They focused on a wide range of simple 1-population and 2-population models with bottleneck, growth, migration and a combination of these parameters. They found that population changes such as ancient temporary bottlenecks would not be inferred correctly regardless of the number of markers available. This set of studies provides valuable information about the use of genomic data in ABC. Our aim is to extend this knowledge by directly comparing ABC results from molecular markers obtained with different types of RRL sequencing techniques, different sequencing effort allocations, and different levels of genomic knowledge. This will hopefully help future ABC users who do not have access to complete genomic data to select methods and develop genomic datasets that are best suited to answer the demographic questions they are addressing.

### General model and datasets

Here, we focused on estimating parameters for a set of 2-population models of demic expansion that are applicable to studies of species invasion, reintroduction, or natural colonization. We tested the power of ABC on these models using a range of marker sets obtainable by RRL methods: datasets with a large number of short genomic reads would correspond to single-end GBS sequencing, whereas fewer but longer diploid sequences correspond to a targeted enrichment approach. For each type of dataset, we quantified the potential benefits of knowing the gametic phase of sequence markers by including or excluding linkage-related statistics at the data-summarizing step. We expect to observe an improvement in the inference for datasets with long sequences. For each model assessed, we also tested the effect of time since colonization. We hypothesize that recent events might be inferred more accurately with datasets containing linkage information, due to the generally higher rate of recombination compared to mutation, and to the potential information contained in long haplotypes. This part of the analysis is also motivated by the fact that overestimates of divergence times are a common result of demographic inference in empirical studies (Holliday et al., 2010) and this upward bias has been found for some demographic scenarios in simulation studies (Benazzo et al., 2015). We therefore aim to explore this potential bias by testing increasingly old events within the same models. As NGS techniques require a trade-off between sample size and individual sequencing depth, and are characterized by high genotyping errors, we explore the effect of different trade-offs at different sequencing error rates. Fumagalli (2013) found that increasing sample size at the cost of decreasing depth was beneficial in the inference of diversity measures and population structure. Here, we extend this hypothesis to ABC inference. Finally, we compared our ABC results with those obtained from an approximate likelihood method using the site frequency spectrum from simulated reduced-representation libraries. As they provide millions of genome-wide SNPs without ascertainment bias, restriction enzyme-based genomic sequencing techniques seem to be particularly well suited to SFS-based inference methods. Comparing SFS results with ABC results on a range of models and datasets will inform future work on demographic inference in non-model organisms.

## Methods

### Demographic models

We focused on a basic 2-population model of demic expansion (fig.1a). A pre-existing population, population 1, is of constant size N_1_. At time T_EXP_ before present, the spatial population expansion begins: population 2 is created by 2 migrants from population 1. Population 2 then grows exponentially between times t=T_EXP_ and t=0 (the present) to size N_2_ at t=0. The rate of population growth *r* is defined by the other parameter values through the formula 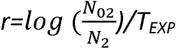. Model 1 therefore has just 3 independent unknown parameters: N_1_, N_2_, and T_EXP_. We created additional models of increasing complexity by adding parameters. In models 2 and 4, the number of founders of population 2, N_02_, is unknown (fig.1b and fig.1d); in models 3 and 4, migration is allowed from population 1 to population 2, with the parameter m_21_ describing a per-generation migration rate (fig.1c and fig.1d). In all four models described above, the mutation rate and the recombination rate are fixed. We chose wide and uniform parameter priors for population sizes to accommodate a wide range of types of organisms, and a log-uniform prior for the timing of the expansion event, as this study intends to focus on more recent rather than ancient expansion events (Table 1).

**Figure 1.**
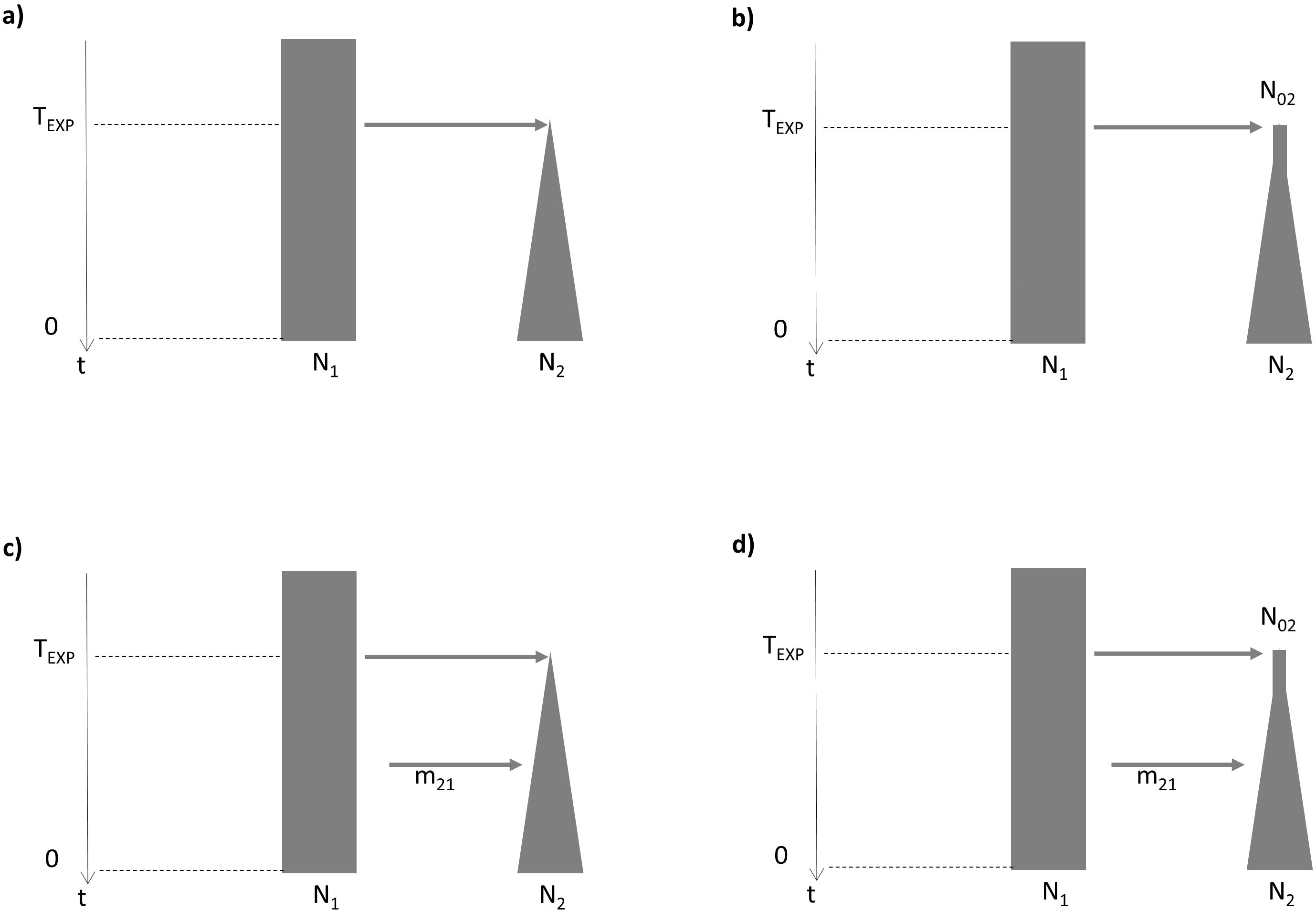
Demographic models. a) Model 1: A three-parameter model of expansion featuring colonization of new population 2 by 2 diploid individuals from population 1 at time T_EXP_. Population 1 is of constant size N_1_, whereas population 2 grows exponentially to size N_2_, its size at present. b) Model 2: the number of founders of population 2 is a variable parameter. c) Model 3: a per-generation migration rate from population 1 to population 2 is added as a parameter. d) Model 4 includes all 5 parameters: N_1_, N_2_, T_EXP_, N_02_, and m_21_.

**Table 1.** Model parameters with their associated prior ranges

| estimated<br>in models | Parameter | Symbol | Prior range | Unit |
| --- | --- | --- | --- | --- |
| - | Mutation rate | $\mu$ | 0.000000009 | - |
| - | recombination rate | R | 0.00000001 | - |
| 1,2,3,4 | population size 1 | $N_1$ | U(10,000:100,000) | ind. |
| 1,2,3,4 | population size 2 | $N_2$ | U(10,000:100,000) | ind. |
| 1,2,3,4 | time of expansion | $T_{EXP}$ | logU(2:10,000) | gen |
| 2,4 | initial population size 2 | $N_{02}$ | U(2:1000) | ind. |
| 3,4 | migration rate from 1 to 2 | $m_{21}$ | U(0.001:0.01) | - |

### Generating sets of coalescent simulations

For each of the four models, we created a set of 1 million simulations with each of the five types of datasets described below, with a fixed number of 10 diploid individuals sampled per population. For datasets corresponding to single-end RADseq sequencing techniques, we simulated 10,000 independent DNA sequences of 100bp each. For datasets corresponding to sequence capture methods, we created 100 independent DNA sequences of 10kb each. Additionally, we explored a range of possible configurations between these two types of datasets (Table 2). With 4 models and 5 types of datasets, we obtained a total of 20 combinations of models and datasets, each with a million simulations. We used the program scrm (Staab et al., 2015), which simulates datasets by creating the ancestral recombination graph following the Wiuf and Hein method (1999). We used custom Rscripts (R Core Team, 2016) inspired by scripts from Shafer et al. (2015) to compute the simulations, and made them available in the supporting information.

**Table 2.** Description of the 5 types of simulated datasets

|  | number of<br>sequences | sequence<br>length<br>(bp) | number of<br>diploid<br>individuals |
| --- | --- | --- | --- |
| 1 | 10,000 | 100 | 20 |
| 2 | 5,000 | 200 | 20 |
| 3 | 1,000 | 1,000 | 20 |
| 4 | 500 | 2,000 | 20 |
| 5 | 100 | 10,000 | 20 |

### Summary statistics

For each simulation we computed all summary statistics available in the program msABC (Pavlidis et al., 2010). The available statistics include diversity statistics (number of segregating sites and θ estimates) and summaries of the SFS (Tajima’s D and Fay and Wu’s H). These statistics were calculated for each population and for the whole sample. The available statistics also include summaries of the 2d-SFS: differentiation measures such as the pairwise F_ST_ and the number of private and shared polymorphisms. Finally the Thomson estimator of T_MRCA_ and its variance were calculated for each population and for the whole sample. To test the effect that knowing haplotype information has on inference, the ABC analysis was performed twice on each model-dataset type combination. The first time, we summarized data using only the statistics mentioned above, which are calculated at the SNP level and therefore are available when the gametic phase of the diploid sequences is unknown. The second inference was performed on the same dataset, but additional statistics Zns (Kelly, 1997), dvk and dvh (Depaulis & Veuille, 1998) based on linkage information were used to summarize the data. These additional statistics are calculated at the haplotype level and so are only available in cases where the gametic phase of the diploid sequences is known. For each set of simulations, we computed the mean and variance of every statistic over all sequence markers in the dataset. As a result, 58 statistics were computed for datasets with known gametic phases (hereafter referred to as “phased”, or “hap.phase 1”), and 43 statistics were computed for datasets with unknown gametic phases (hereafter referred to as “unphased”, or “hap.phase 0”).

Using a high number of statistics to summarize genetic data has harmful effects on the quality of the ABC inference, a problem commonly referred to as the “curse of dimensionality” (Blum et al., 2013). We used the partial least squares (PLS) method implemented in ABCtoolbox (Wegmann et al., 2010) to reduce the number of statistics to 5-7 PLS components (see Supplemental methods for details).

### Pseudo-observed datasets

For each set of 1M simulations, we created a corresponding set of 100 pseudo observed datasets (PODs), with parameters randomly chosen from the same priors as for the set of 1M simulations. By doing so we assume that priors are reliable and reflect the true, unknown distribution of the PODs. These were then summarized with the same summary statistics as their corresponding set of 1M simulations.

### ABC estimation

We performed the ABC estimation using each POD as the observed dataset to obtain parameter estimates. The standard ESTIMATE algorithm from the program ABCtoolbox (Wegmann et al., 2010) was used for all ABC computations to create posterior probabilities from the corresponding set of 1M simulations, with a post-sampling regression adjustment through ABC-GLM (Leuenberger & Wegmann, 2010). We fixed the tolerance parameter to 10^−3^, a compromise between having a tolerance threshold value as low as possible (Li & Jakobsson, 2012) and keeping an appropriate number of simulations to estimate the posterior from.

### Validation

For each combination of model and type of dataset, we computed a measure of precision and accuracy called the relative prediction error (RPE), the ratio of the mean squared error over the variance of the prior, which follows equation (2):

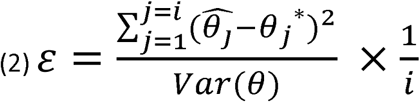

where *Var*(*θ*) is the variance of the prior distribution and i is the number of observations. The RPE was computed on 1,000 PODs. The advantage of using RPE as a validation statistic is that it directly indicates the contribution of the genetic dataset to the estimation of the posterior. Another attractive feature of the RPE is that it allows comparisons between parameters, as it scales from 0 (precise estimate) to 1 and beyond (in the case of a consistent bias in estimation).

As an additional measure of precision, the 95% highest posterior density interval (HDI) was calculated on a set of 100 PODs for each combination of model and dataset type. This measure is defined as the shortest continuous interval with an integrated posterior density of a certain value (Wegmann et al., 2010). For each combination of model and dataset type we reported the 95% HDI coverage, i.e. the number of times (out of 100) the true parameter value fell within the 95% HDI, expecting values close to 95.

### Testing the effect of T_EXP_ on parameter estimation

To test the effect of the time of expansion on the precision of the ABC estimation, we created 100 PODs for each set of 1M simulations and 12 fixed values of logT_Exp_ spanning the prior range. RPE and 95%HDI were calculated from the results of each set of 100 PODs.

### Effect of sequencing effort allocation and sequencing error

The main challenge when developing genomic markers is managing sequencing and variant calling errors. Sequencing a large number of individuals might increase the precision of population genetics inference, but with a fixed sequencing budget, this comes at the cost of reduced individual sequencing depth, which in turn can affect variant calling and estimation of allelic frequencies (Fumagalli, 2013). We explored this challenge focusing on model 2 and dataset type 2. We chose a realistic fixed sequencing effort and derived 3 fixed sampling strategies from it: 250 sampled individuals at a mean individual depth of 4, 100 individuals with depth 10, and 20 individuals with depth 50. We then incorporated three per-nucleotide sequencing error rates (0, 10^−2^, 10^−3^), and applied them to each category described above. The resulting 9 categories of PODs, as well as “perfect” datasets (no depth sampling and no error) were all simulated using the same 10 parameter combinations. Further details about the creation of “imperfect” PODs can be found in the supplemental methods. Once these imperfect PODs were created and summarized, ABC was performed to estimate their true parameter values. Two additional sets of 1M simulations needed to be created to match the number of individuals sampled per population: one with 100 diploids per population, and the second with 250. It the latter case, we only created 610,000 simulations because of computation time limitations. The same tolerance (0.001) as all other runs was used for the estimation.

### Comparing ABC and SFS estimation

We simulated 10,000 independent DNA sequences of 100bp each for the 4 demographic models 10 times. The resulting 40 datasets were input into both ABCtoolbox and fastsimcoal2, which uses the SFS to approximate a composite likelihood from a large number of simulations through a conditional maximization algorithm (see supplemental methods). We compared the results from the two methods using RPE, credible intervals and confidence intervals.

## Results

A total of 20 combinations of models and datasets were used as input for ABC simulations (Tables 1 and 2), resulting in a total of 20 million simulated datasets available for analysis, training simulation sets and PODs. Each set of 1M simulations was used in two runs of estimation: one including all summary statistics available in msABC, the other one excluding statistics based on linkage information, for a total of 40 ABC estimations.

### Effect of model complexity on the precision of parameter estimates

In general, the ability to infer demographic history declined rapidly as model complexity increased. The simplest model (1), estimating only population sizes N_1_ and N_2_ and the log-transformed time of expansion T_EXP_, allowed the expansion event to be dated accurately. Models 2 and 3 each had 4 parameters: model 2 included the number of founders N_02_ and model 3 allowed migration from population 1 to population 2 (m_21_). For both model 2 and 3, logT_EXP_ was inferred with slightly lower precision than for model 1. Finally, scenarios corresponding to model 4, which had all 5 parameters, failed to be correctly inferred.

Not all parameter estimates were sensitive to the addition of parameters in the models: the precision of contemporary population size estimates N_1_ and N_2_ were independent of model complexity. RPE values for N_1_, which was constant over generations, were mostly below 0.05 for the four models assessed (fig. 2). The 95% highest posterior density intervals ranged from 3,000 to 60,000. For N_2_, the contemporary population 2 size after exponential growth, 95% HDI intervals were about as wide as the prior range, indicating a failure to estimate this parameter in all four models (fig. 3).

**Figure 2.**
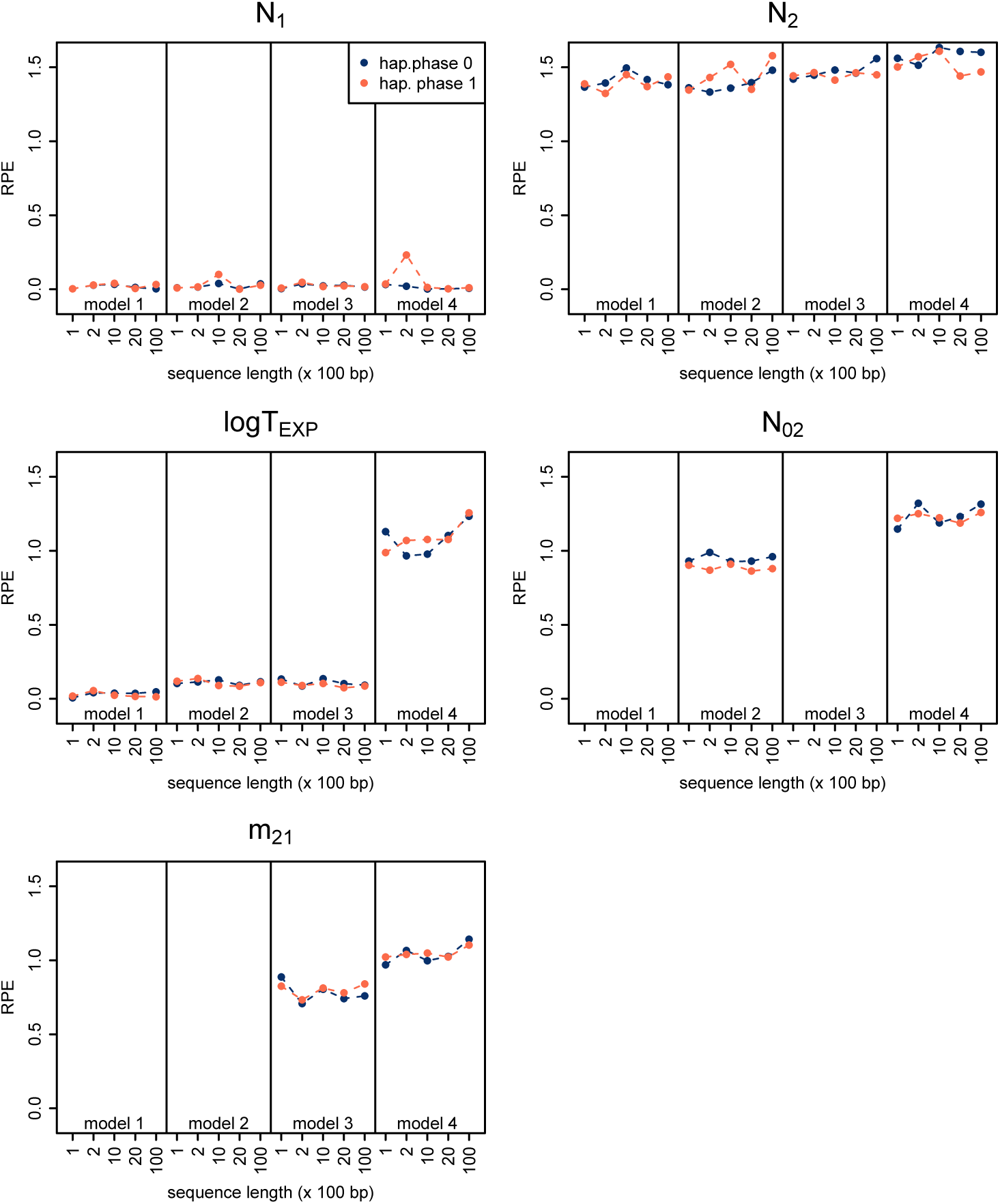
Relative prediction error (RPE) calculated from the results of ABC analyses of 20 different combinations of demographic models and sampling designs (x-axis). For each combination, ABC was performed on simulated datasets summarized with statistics including linkage-based measures (hap. phase 1) and on the same set of simulations summarized with only SNP-based statistics (hap. phase 0). RPE values were calculated from the ABC estimation results of 1000 datasets with parameter values randomly drawn from their prior distributions.

**Figure 3.**
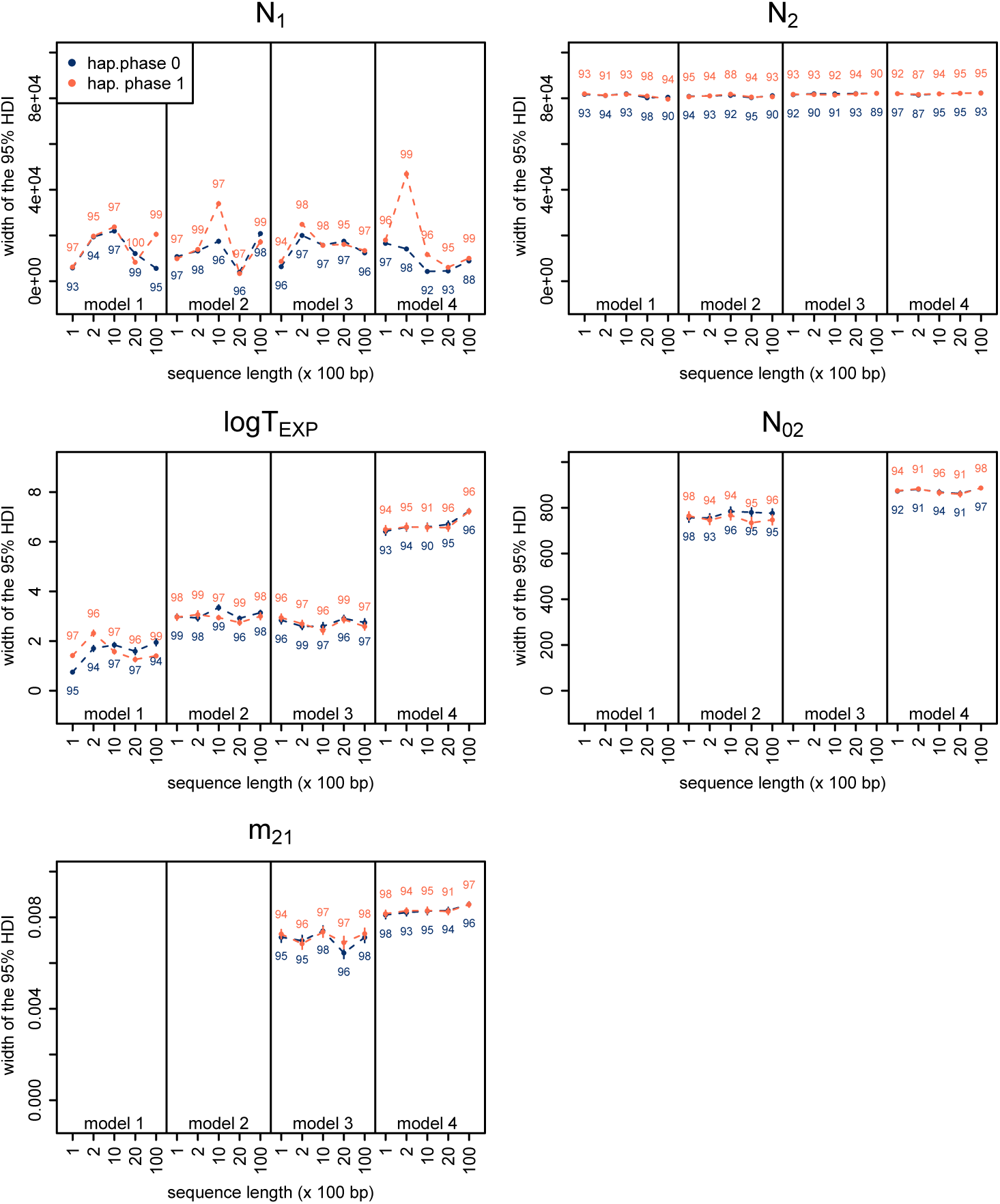
Width of the 95% highest posterior density intervals calculated from the results of ABC analyses of 20 different combinations of demographic models and sampling designs. Error bars represent standard errors (N=100 PODs). See caption of figure 2 for more details.

The expansion time T_EXP_ was generally well estimated in model 1, which is the simplest 3-parameter model (fig. 2) with no migration between demes and the number of founders set to 2. For this model, the RPE was mostly below 0.1. The precision of logT_EXP_ estimation was almost as high for the two 4-parameter models, where the number of founders N_02_ (model 2) is unknown and needs to be estimated, or where migration from population 1 to population 2 is likely (model 3). For these two models, the RPE is below 0.2. The ABC analysis of the 5-parameter model (model 4) was unable to recover the true T_EXP_ value.

Estimates of the number of founders of population 2 (N_02_) and migration rate from population 1 to 2 (m_21_) were surprisingly imprecise in models of low complexity (model 2 and 3) and could not be recovered at all in model 4 (fig.2 and 3).

Models 1 to 4 all rely on population 2 growing exponentially from T_EXP_ to the present time. We tested whether demographic parameters could be estimated more successfully in a model where population 2 goes through a single sudden population change instead of exponential growth. We created a new set of 1M simulations based on model 2 (where N_02_ is a varying parameter) and dataset type 1 (many short sequences) and a smaller prior range for T_EXP_ (2-500 generations). In the new model the size of population 2 changes from N_02_ to N_2_ at T_EXP_/10 and remains constant before and after T_EXP_/10. These modifications brought no improvements to any of the parameter estimates (Table S1).

### Do sequence length and linkage-related statistics improve the estimation?

The addition of linkage statistics available in msABC brought no notable improvement in the RPE and 95% HDI of parameter estimates for all models (fig.2 and fig.3). It even seems to make the estimation of N_1_ less precise in some cases for model 1, 2 and 4, although this pattern is inconsistent across dataset types. ABC performance on models 3 and 4 seemed to be slightly more dependent on sequence length, with the inference on large sequences marginally benefitting from haplotype information.

### Quality of parameter estimates across prior ranges

For each parameter, we visualized estimated values and 95% HDI of ABC results in relation to true parameter values to assess performance over the prior range. Results for the 3-parameter model (model 1) and dataset types 1 and 5 are shown in fig. 4a and 4b, respectively. Results for the complete set of models are available in supplemental fig. S1. Consistently across models, estimates of N_2_, N_02_, and m_21_ are largely inaccurate regardless of the true value, with HDI ranges as wide as the prior range. Conversely, N_1_ estimates are accurate in all models regardless of the true N_1_ value. Unlike N_1_, the values of T_EXP_ have an impact on the precision of their respective estimates. Accuracy and precision of T_EXP_ estimates for models 1 and 3 decrease with increasing true value. Interestingly, the opposite pattern is observed for model 2: more recent events are less precisely inferred than ancient ones (fig. S1, pp. 11-20). Results for model 4 show a “cross” pattern where most PODs’ logT_EXP_ values are correctly estimated but some PODs with extreme logT_EXP_ values show estimates at the opposite extreme (fig. S1, pp. 31-32). This pattern suggests a complex multivariate relationship between model parameters and statistics.

**Figure 4.**
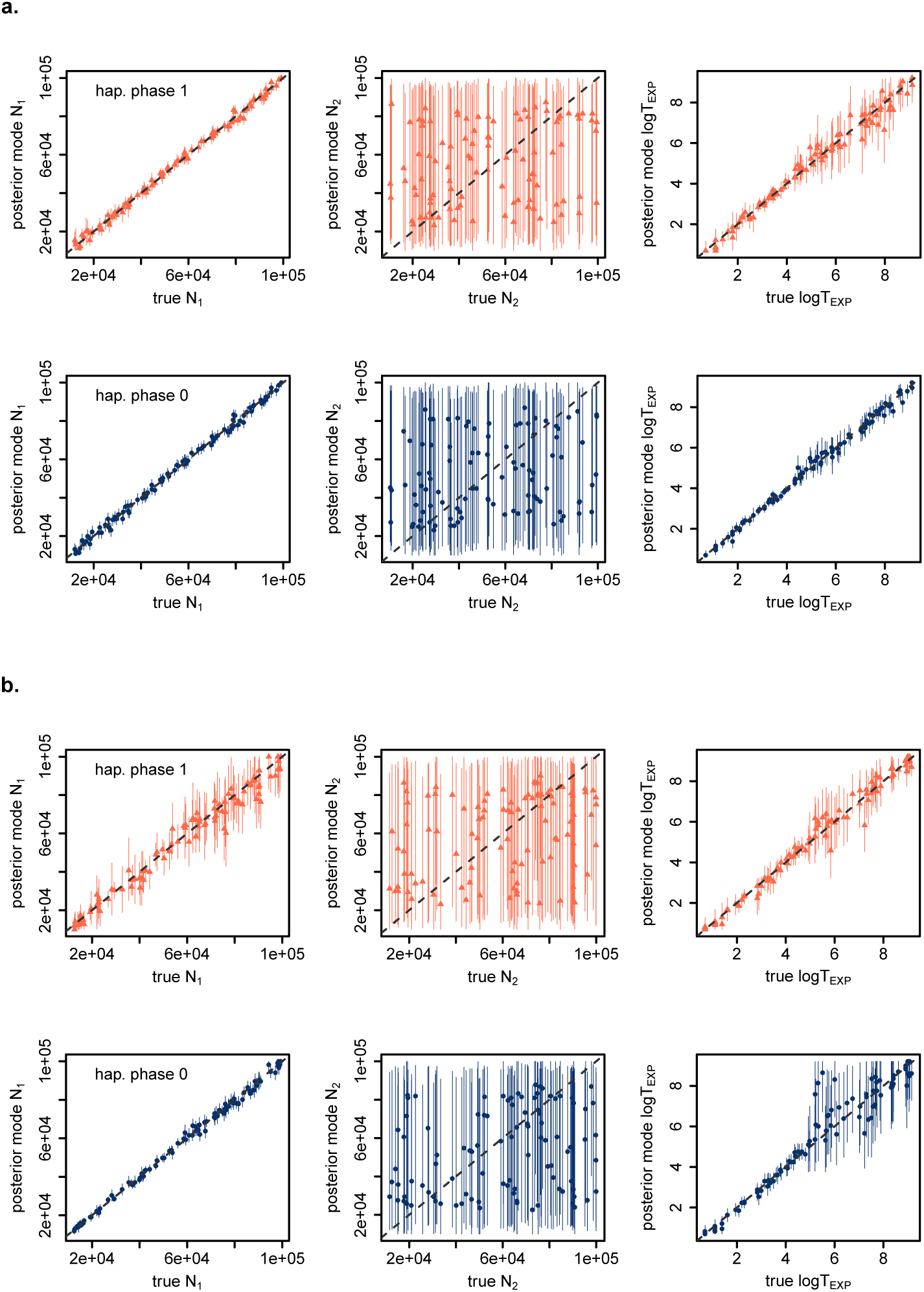
Accuracy of parameter estimates for model 1. Within a plot, each datapoint corresponds to the estimated value of the parameter (mode of the posterior) vs. the true parameter value for one POD. Results are shown for a total of 100 PODs. Error bars correspond to the 95% HDI around the estimate. a) Results with datasets of type 1 (10,000 sequences of 100bp). Top panel shows results on unphased datasets, bottom panel shows results for phased datasets. b) Results with datasets of type 5 (100 sequences of 10,000bp). Top panel shows results on unphased datasets, bottom panel shows results for phased datasets.

### Effect of the time of the expansion event on the estimation

We tested whether older expansion events are generally more difficult to characterize than recent ones within the time range specified by the prior. To do this, we studied the effect of the true T_EXP_ value on the precision of parameter estimates. We find different trends among the 4 models (fig. 5, S2, and S3). The precision of inference on model 1 is higher at low T_EXP_ values and decreases at logT_EXP_>4. Conversely, for model 2, older events are generally better inferred: estimates of T_EXP_ and N_02_ increase in precision as T_EXP_ increases, as shown by the RPE (fig. S2, p.2) and the 95% HDI (fig. S3, p.2). Model 3 shows the best results for moderately recent expansion events (3 < logT_EXP_ < 4), as shown by RPE and 95% HDI of T_EXP_ and m_21_ (fig.S2 and S3). Finally, results for model 4 show high values of RPE and 95% HDI for all parameters, with RPE values mostly above 0.5.

**Figure 5.**
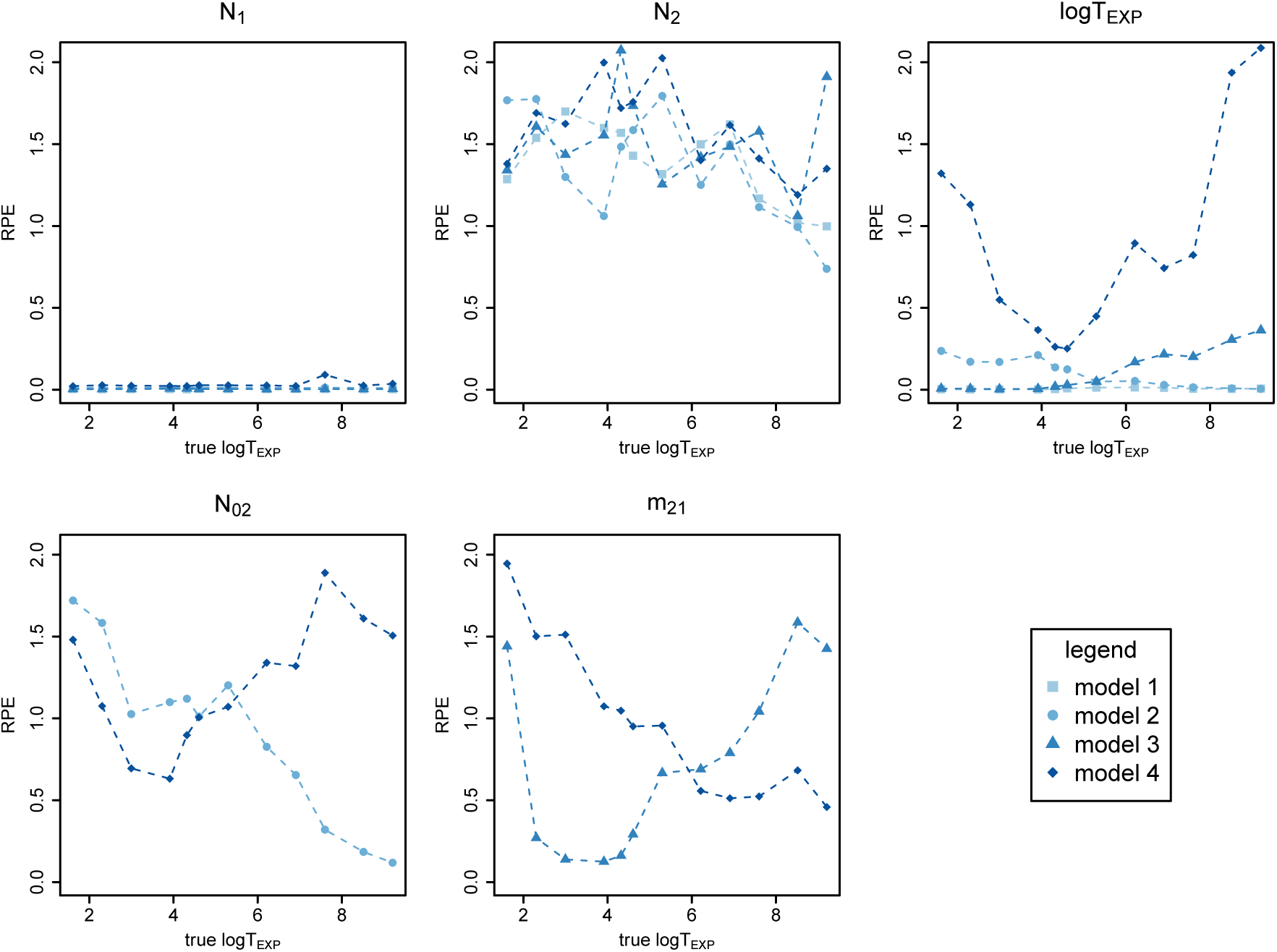
RPE of model parameters for different fixed values of T_EXP_. Results are shown for ABC runs with datasets of type 1 (10k sequences, 100-bp long). For a given parameter, results from different models are shown in the same plot window with different characters and colours. To see results for other model-dataset combinations as well as 95% HDI results, please see supporting information.

### Effect of sequencing effort allocation and sequencing error

Focusing on model 2 and datasets of 5,000 × 200bp sequences, we simulated sequencing and variant calling for three different sample size and depth combinations. The RPE of parameter estimates for 13 tested PODs is represented in fig. 6. Depth of sequencing (dp) has very little effect on the precision of estimates: only N_1_ and logT_EXP_ have a marginally higher RPE when sequencing depth is simulated. Error rates affect N_02_ estimates at low depth (N=250, dp=250), as well as logT_EXP_ estimates at low sample size (N=20, dp=50). The estimation is otherwise robust to introduced errors. For a given set of PODs (e.g. N=250, dp=4), the precision lost in a parameter estimate because of an error rate of 0.01 (N_02_) is gained on another parameter (N_1_), reflecting the limitations of the model estimation process rather than the effect of sequencing error. However, the results suggest that choosing a larger sample size with a shallower individual sequencing depth improves estimation over other strategies, especially for the estimation of logT_EXP_.

**Figure 6.**
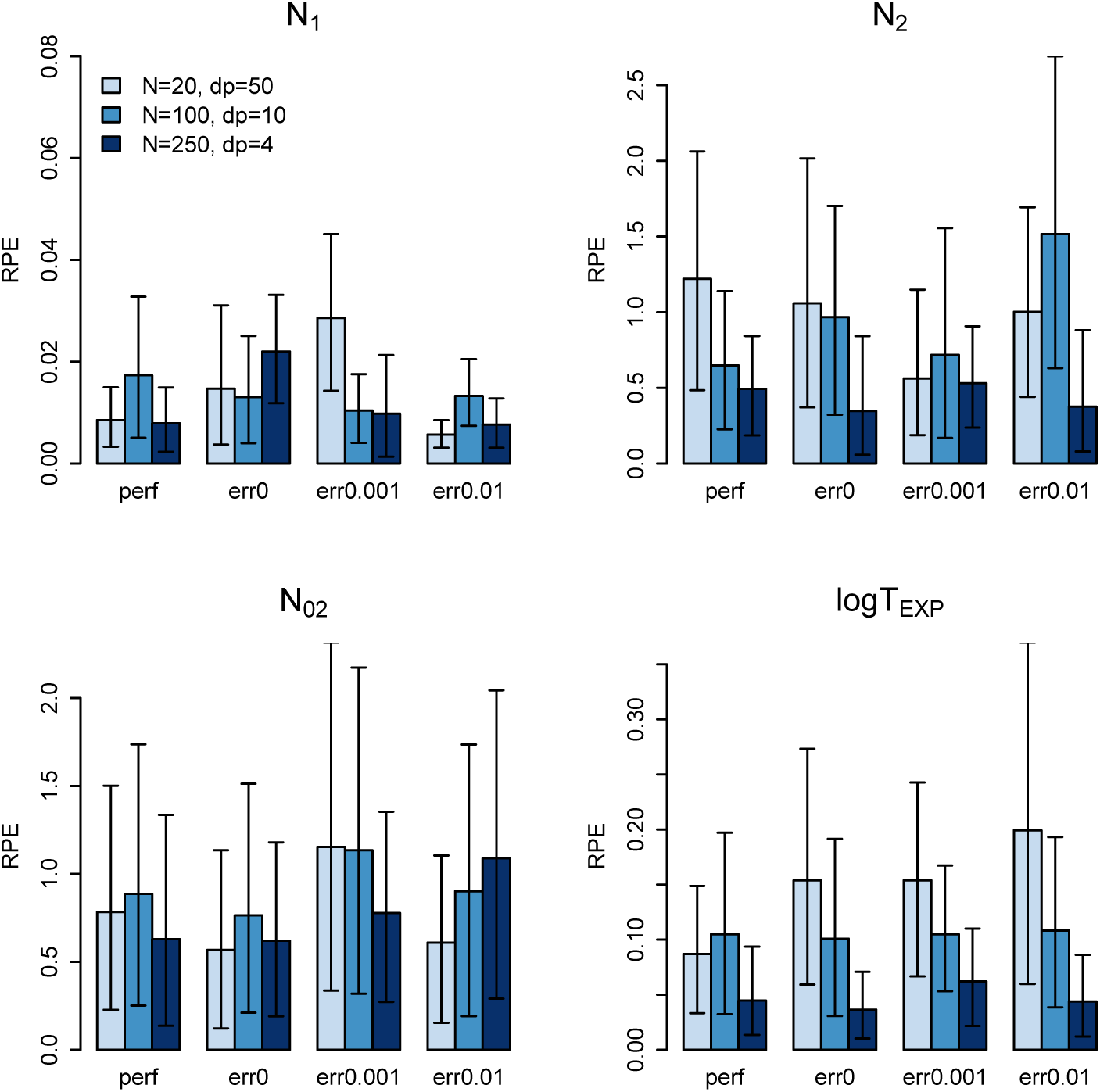
RPE and bootstrapped confidence intervals of model 2 parameters under different sequencing strategies and per-nucleotide error rates. N corresponds to the number of diploid individuals sequenced, dp to the mean individual sequencing depth. “perf” corresponds to perfect datasets whereas “err0”, “err0.001” and “err0.01” correspond to datasets where the sequencing process was simulated, with depth sampling and errors introduced at rates 0, 0.001, and 0.01 substitutions per nucleotide respectively. 13 PODs were used for each treatment.

### Comparing ABC with SFS estimation using an approximate composite likelihood

Figures 7 and 8 illustrate the performance of ABC and approximate composite likelihood from the SFS for all models performed with datasets of 10,000 100-kb sequences. Both methods gave similar results in terms of precision of parameter estimates. The SFS-based method performed slightly better than ABC in the model with migration (model 3), but the precision of ABC estimates was superior for model 2 (fig.7). The approximate composite likelihood method generally provided narrower 95% confidence intervals (fig.8).

**Figure 7.**
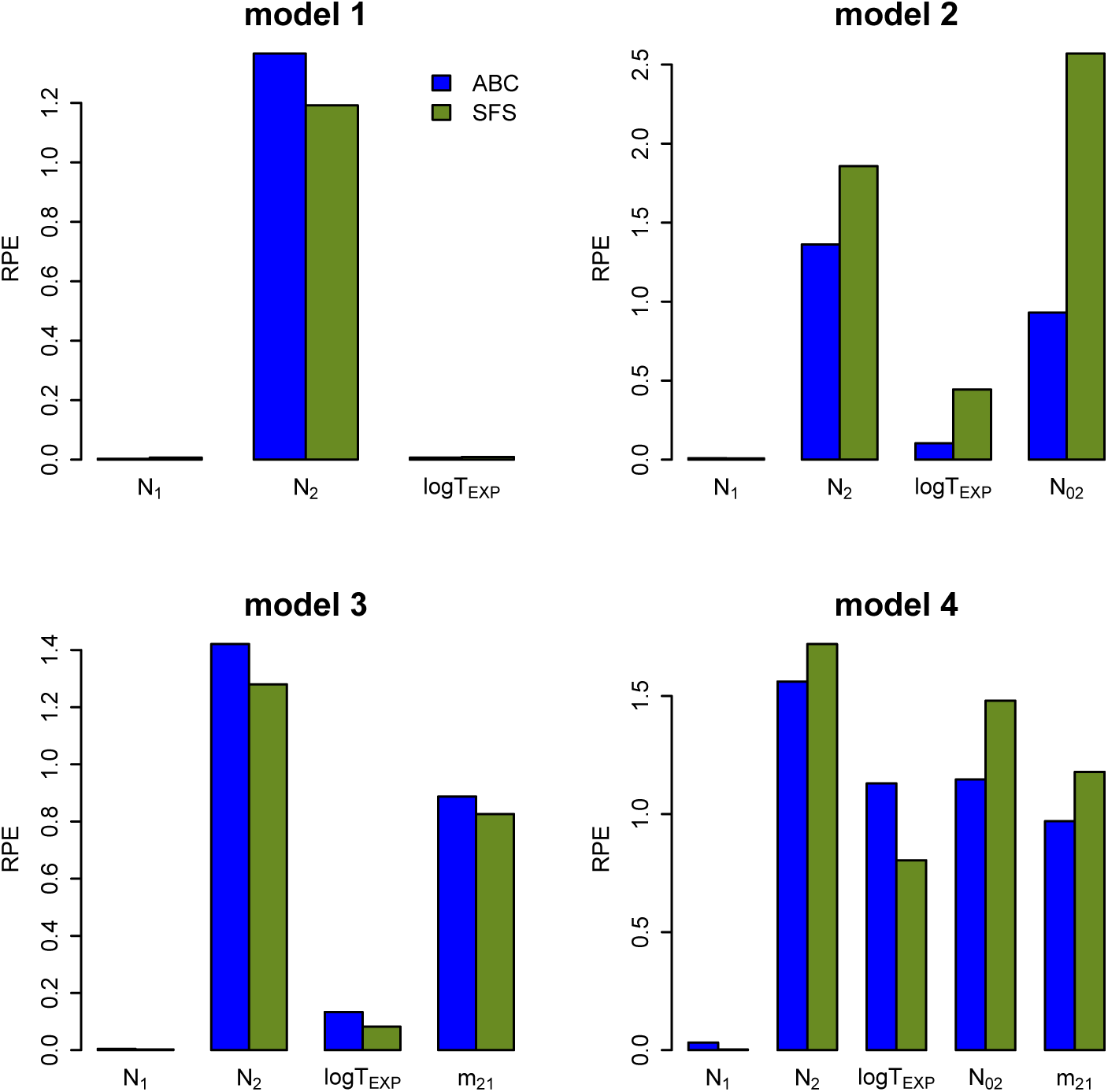
RPE calculated from 100 datasets for models 1 to 4 using two different inference methods: ABC, computed on SNP-level summary statistics, and approximate composite likelihood, computed from the SFS. In both cases, datasets had 10,000 sequences of 100bp genotyped in 20 diploid individuals.

**Figure 8.**
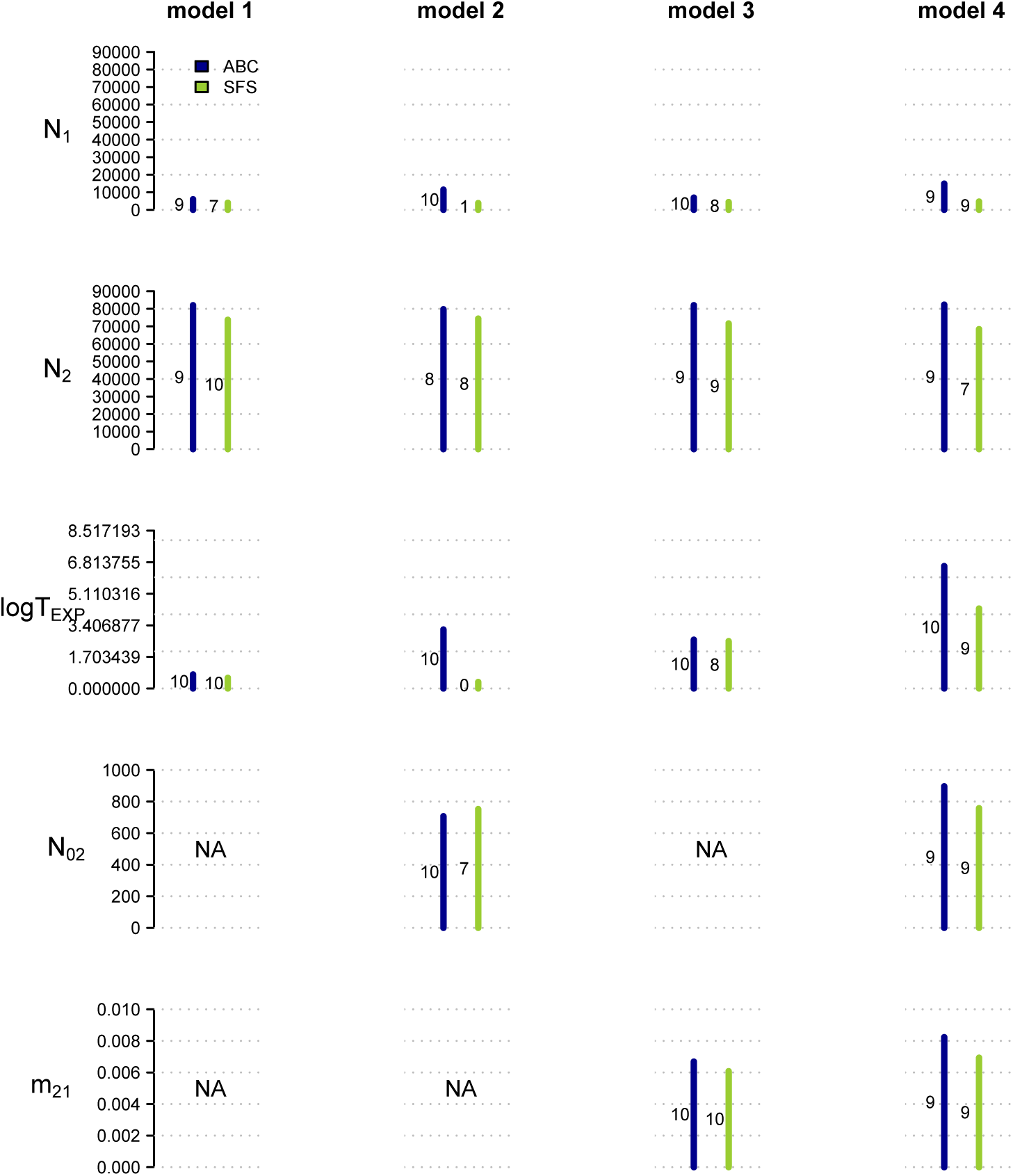
Width of the 95% HDI from ABC results, compared to 95% CI from the SFS inference method. For each of the four demographic models, the same 10 simulated datasets were used as pseudo-observed datasets for both the ABC and the SFS runs. HDI and CI widths were calculated from 100 bootstraps. Numbers correspond to the coverage of 95% CI (out of 10 PODs). PODs had 10,000 sequences of 100bp genotyped in 20 diploid individuals.

## Discussion

We explored the ability of approximate Bayesian computation to characterize a recent event of spatial expansion from one population of constant size to a new and growing population, a model which can be broadly applied to studies of species range expansion, invasion biology, or reintroduction of endangered species. We found that regardless of model complexity, estimates of the size of the growing, newly founded population (N_2_) are poor. However this did not prevent successful estimation of other parameters (N_1_, logT_EXP_, and in restricted cases N_02_). Failure to estimate N_2_ does not come as a surprise: estimates of past changes in effective population size from one punctual sampling event commonly rely on linkage information between markers, a calculation not readily available in ABC packages (Beaumont, 2003). Our result that models of higher complexity are harder to estimate was expected, but in the case of our expansion models, this trend leads surprisingly quickly to a complete failure to estimate any parameter, as soon as 5 parameters are involved. While expansion timing was precisely estimated in the 3- and 4- parameter models, it could not be recovered in the 5-parameter model. ABC on model 2, the 4-parameter model including the number of founders but no subsequent migration, successfully estimated all parameters (except N_2_) for old expansion events. In contrast, for model 3, the 4-parameter model including migration between demes, estimations were more successful for recent events. These results highlight the potential importance of taking into account the timing of an expansion event when predicting estimation success for a given demographic model. The difficulty of estimating the time of a founding event with subsequent migration was also reported by Robinson et al. (2014); however, we show here that for a moderately recent event (10 to 100 generations), it is possible.

### Implications of including haplotype information

Analyses based on unphased sequences exploring similar models to those used here have shown encouraging results (Robinson et al., 2014). However, no study to date has explicitly compared datasets of phased and unphased sequences using the same models and same amount of data. Here, we quantified the benefits of using phased haplotype sequences over single SNPs by including or leaving out LD-based and haplotype-level statistics at the data summarization step of the ABC inference. Surprisingly, haplotype information did not substantially improve the precision of parameter estimates, even when 10-kb sequences were used as markers. Li and Jakobsson (2012) explored ABC with similar 2-population split models and a similar fixed population-wise per-generation recombination rate as in our study. When they tested different combinations of summary statistics, their results did not demonstrate any obvious superiority of LD-based statistics over SNP-based statistics. They concluded that the selected summary statistics should capture as many different aspects of the data as possible, with as little redundancy as possible. Potentially, phasing the data may not have improved inferences because the extent of linkage that the chosen statistics are sensitive to differs from the linkage actually present in the simulated data. Future work when dealing with phased data would require developing expectations of LD levels and creating or choosing statistics that cover the extent of LD likely to be present in the data.

One needs to be aware of the difficulties associated with the use of LD information. Firstly, ABC on phased data requires reasonable knowledge of recombination rates and variability across the genome. The recombination rate needs to be included as a parameter along with demographic parameters, or as a nuisance parameter with a hyper-prior. Secondly, simulating the coalescent with recombination is a complicated process and comes at high computational costs (McVean & Cardin, 2005). With high recombination rates or very long sequences, coalescent simulations might take so long to run that one would instead use a more efficient inference method than ABC. Moreover, translating genome-wide observed data into a set of summary statistic values that are readily useable by ABC programs and comparable to simulated datasets can be a challenge. File input formats in most programs are currently not compatible with sequence information, and many summary statistics programs do not offer haplotype-level calculations. Thirdly, when aligning reads to a fragmented and incomplete reference genome, as is often the case for non-model organisms, defining haplotypes can be tricky. One also needs to address problems of sequencing errors, paralogous sequences and imperfect mapping. Inevitable sequencing uncertainties will affect haplotype statistics more strongly than single-SNP diversity measures. Data processing errors and filters can severely bias inferences, to the extent of supporting the wrong demographic model, as revealed by Shafer et al. (2016). Finally, targeted sequence capture will result in thousands of markers of various lengths. Setting up simulations that correspond closely to an observed dataset requires approximating the distribution of sequence lengths, and this may also affect inferences, especially if variances of summary statistics are included at the data summarization step. Considering the difficulty of obtaining reliable haplotype information in non-model organisms, the potential difficulties of adapting the use of long sequences to currently available ABC programs, and computational time, our results tend to suggest that using SNP-level information from GBS-type data is preferable over targeted sequence capture.

### Choosing summary statistics

It is important to note that all the results presented here are only valid in the context of our choice of summary statistics. In the present study, we decided to use the first and second moment of all statistics available in msABC, and to reduce the dimensionality with a PLS transformation. Several previous publications have performed simulations either using the two first moments of summary statistics (Li & Jakobsson, 2012) or only using the mean (Shafer et al., 2015). To our knowledge, only Robinson et al. (2014) tested the use of 4 moments for summary statistics for models of divergence with admixture. They compared their results with those obtained using only the mean and found that the mean alone was sufficient. Although the two first moments may not be the most representative summaries for some statistics, adding higher-level moments will come at a computational cost.

It is widely recognized that choosing a set of summary statistics is probably the most challenging step for ABC users. For instance, the optimal set of statistics for parameter estimation in a given model might differ from the optimal set of statistics to discriminate between demographic models. As insufficient summary statistics have detrimental effects on model selection (Robert et al., 2011), Fernhead and Prangle (2012) introduced “semi-automatic ABC”, which relies on an ABC pilot run and a subsequent linear regression to choose the most appropriate set of summary statistics. Similarly, ABCtoolbox 2.0 implements a statistical selection step based on the incremental assessment of inference power with the addition of summary statistics. However, documentation is lacking for this new feature of the program. These improvements constitute a promising step towards a more rigourous statistical framework for the automatic selection of ABC summary statistics.

### Sequencing effort: go large and shallow!

We found that “imperfect” datasets created with a high number of individuals sequenced at a low individual depth seemed to perform consistently better for most parameters than datasets with fewer individuals and higher depth. This is consistent with Fumagalli (2013), who studied the same trade-offs on diversity statistics under various demographic settings. This result seems to hold even with simulations with moderate or high sequencing error rates, although this is difficult to conclude with confidence considering the large bootstrapped confidence intervals (fig.6). It is worth noting that if the error rate is not properly estimated during the genotype calling process, more errors will be present in the final dataset and it is likely that ABC results will be impacted for all sequencing strategies, especially those with low depth. As ABC summary statistics rely on the SFS and not on individual genotypes, we suggest that future ABC users sequence large sample sizes at low depth. In this case, estimating the SFS or derived statistics following methods such as described in Nielsen et al. (2012) and Fumagalli et al. (2014) has proven more successful than genotype calling in inferring the SFS. There is unfortunately no straightforward program or pipeline of compatible programs incorporating these methods into an ABC framework. One possibility is to summarize the SFS into quantiles and to use the latter as summary statistics in a classic ABC run. Such a process would need to be further tested.

### Comparing ABC to other methods

We did not find large differences in the precision of parameter estimates between ABC and the SFS-based likelihood method implemented in fastsimcoal2. Shafer et al. (2015) found a similar result while comparing the performance of ABC with a SFS-based inference implemented in *δαδi* (Gutenkunst et al., 2009). They found that *δαδi* tends to overestimate the time of population split and bottleneck events, a trend not supported by our findings with *fastsimcoal.* In addition to parameter estimation, Shafer *et al.* (2015) tested the performance of both methods for model selection and found ABC more accurate, especially in the case of bottleneck scenarios.

ABC has proven moderately useful for demographic inference with long, genome-wide haplotypes but comparisons with alternative approaches are scarce. Notable examples include Nadachowska-Brzyska et al. (2013), who used ABC and PSMC in a complementary way. Robinson et al. (2014) compared their ABC results with an exact likelihood method developed by Lohse et al. (2011) and found that ABC resulted in more uncertainty, especially in model comparisons. As ABC performance with linkage information needs to be further explored, comparisons to emerging analytical methods based on whole genomes or long sequences such as MSMC (Schiffels & Durbin, 2014) or identity-by-descent haplotype sharing (Harris & Nielsen, 2013) will greatly help refine methods for demographic inference using data at a genomic scale.

Theoretical improvements of ABC methods are emerging rapidly. Although the results presented here do not show that ABC benefits greatly from the use of deeper genomic datasets, the versatility of ABC might be key to its useful applications in a wide variety of fields, even those progressing rapidly such as population genetics. Constant methodological improvement, however, requires regular updates to available ABC programs.

## Acknowledgements

J.E was supported by an NSERC Discovery Grant to S.N.A and a Strategic Recruitment Fellowship from the Faculty of Forestry, University of British Columbia. We thank Michael Whitlock for his insightful comments on the manuscript, and Daniel Wegmann for providing bioinformatics support. Thanks to the two anonymous reviewers for their helpful comments.

## Data accessibility

All relevant information to reproduce this study is included in this manuscript and supporting information.

## Author contributions

J.S.E and S.N.A conceived the study. J.S.E performed simulations and analysed the data. J.S.E wrote the manuscript with input from from S.N.A.

## Supporting information

Additional supporting information including methods, figures and scripts can be found online.

